# Sensitivity to visual features in inattentional blindness

**DOI:** 10.1101/2024.05.18.593967

**Authors:** Makaela Nartker, Chaz Firestone, Howard Egeth, Ian Phillips

## Abstract

The relation between attention, perception and awareness is among the most fundamental problems in the science of the mind. One of the most striking and well-known phenomena bearing on this question is *inattentional blindness* (IB; Neisser & Becklen, 1975; Mack & Rock, 1998; Most et al., 2001, 2005). In IB, naïve observers fail to report clearly visible stimuli when their attention is otherwise engaged—famously even missing a gorilla parading before their eyes (Simons & Chabris, 1999). This phenomenon and the research programs it has motivated carry tremendous theoretical significance, both as crucial evidence that awareness requires attention (Cohen et al., 2012; Prinz, 2012; Noah & Mangun, 2020) and as a key tool in seeking the neural correlates of consciousness (Rees et al., 1999; Pitts et al., 2014; Hutchinson, 2019). However, these and other implications critically rest on a notoriously biased measure: asking participants whether they noticed anything unusual (and interpreting negative answers as reflecting a complete lack of perception). Here, in the largest ever set of IB studies, we show that, as a group, inattentionally blind participants can successfully report the location, color and shape of the stimuli they deny noticing. This residual visual sensitivity shows that perceptual information remains accessible in IB. We further show that subjective reports in IB are conservative, by introducing absent trials where no IB stimulus is presented; this approach allows us to show for the first time that observers collectively show a systematic bias to report not noticing in IB—essentially ‘playing it safe’ in reporting their sensitivity. This pair of results is consistent with an alternative hypothesis about IB, namely that inattentionally blind subjects retain some degree of awareness of the stimuli they fail to report. Overall, these data provide the strongest evidence to date of significant residual visual sensitivity in IB. They also challenge the use of inattentional blindness to argue that awareness requires attention.

## Main

One of the most pervasive and compelling intuitions about perception is that we see what is right in front of us: If a large enough stimulus were to appear directly before our eyes (with good lighting, a well-functioning sensory apparatus, no special camouflage, etc.), we would see it and could easily report features such as its color, shape, and location. However, this seemingly secure assumption is challenged by perhaps the best-known result in contemporary perception science: The phenomenon of inattentional blindness (IB). In IB, engaging in an attentionally demanding task (e.g., judging the relative lengths of two briefly presented lines, or counting basketball passes) causes observers to miss large, highly visible, but unexpected stimuli appearing right before their eyes (e.g., a high-contrast novel shape, or a parading gorilla; Figure 1). Whereas in ordinary circumstances these stimuli would be extremely salient and easily noticed, under conditions of inattention we seemingly do not see them at all.

**Figure 1.**
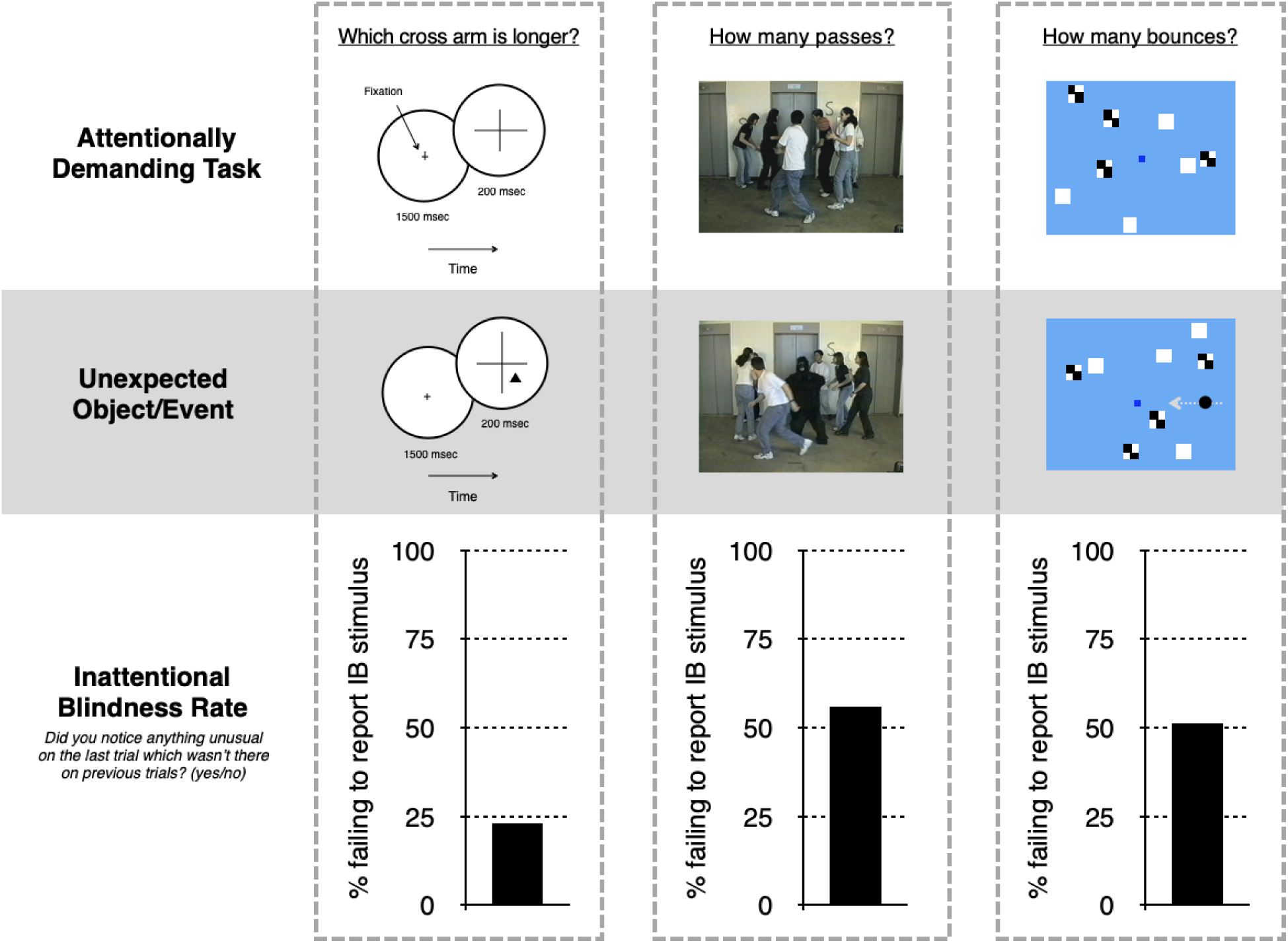
Influential inattentional blindness paradigms and their results. (Left) In a representative study from Mack & Rock (1998), subjects fixated in the center of the circle and reported which arm of a centrally presented cross was longer. On the critical trial, an unexpected shape appeared in the near periphery at the same time as the cross and subjects were asked whether they noticed anything unusual. **(Center)** In Simons & Chabris (1999), subjects counted the number of basketball passes made by individuals wearing white shirts. (Images drawn from Simons and Chabris, 1999, reprinted with permission Dan Simons.) On the critical trial, a woman in a gorilla suit paraded through the display for five seconds, and subjects were asked whether they noticed anything unusual. **(Right)** In Wood & Simons (2017; based on Most et al., 2001), subjects fixated on the blue square and reported the number of times the white or checkerboard squares bounced off the walls of the display. On the critical trial, a black circle entered from the right and crossed the display for several seconds, and subjects were asked whether they noticed anything unusual. In all three experiments (and many others in this literature), a considerable proportion of subjects reported not noticing these unexpected stimuli.

IB is an extremely robust phenomenon, demonstrated in a stunning variety of laboratory and real-world contexts across half a century of research (Neisser & Becklen, 1975; Moore & Egeth, 1997; Mack & Rock, 1998; Simons & Chabris, 1999; Most et al., 2005; Drew et al., 2013; Murphy & Greene, 2016). Although the proportion of subjects demonstrating IB (the “IB rate”) varies widely depending on the protocol, it can be remarkably high (e.g., ∼50% in the case of the famous gorilla, and over 80% in some of Mack & Rock’s studies; for reviews see Simons, 2000; De Brigard & Prinz, 2010; Jensen et al., 2011; Nobre et al., 2020; Redlich et al., 2021).

IB is also an extremely significant phenomenon. According to the “consensus view” (Noah & Mangun, 2020; see also Mack & Rock, 1998; Dehaene et al., 2006; Cohen et al., 2012; Prinz, 2012, 2015; though see Koch & Tsuchiya, 2007; Maier & Tsuchiya, 2020), subjects undergoing IB “have no awareness at all of the stimulus object” (Rock et al., 1992), such that “one can have one’s eyes focused on an object or event … without seeing it at all” (Carruthers, 2015). This interpretation has endowed IB with numerous theoretical and practical implications. First, IB is a key piece of evidence for the more general claim that awareness requires attention (or, conversely, that “without attention, conscious perception cannot occur”; Dehaene et al., 2006), and so has been used to support various leading theories of consciousness on which attention plays a critical role, such as global neuronal workspace theory (Dehaene & Naccache, 2001), attended intermediate representation (AIR) theory (Prinz, 2015), and attention schema theory (Webb & Graziano, 2015). Second, conceived as a tool to abolish awareness, IB is frequently used by neuroscientists to measure neural activity in the absence of consciousness (Rees et al., 2000; Pitts et al., 2014; Hutchinson, 2019), in the hope of isolating the neural correlates of consciousness. Third, IB challenges cherished assumptions about our ability to perceive the world around us—inspiring substantial theoretical and philosophical debate (e.g., O’Regan & Noë, 2001; Lamme, 2003; Block, 2011; Cohen & Dennett, 2011) and rightly contributing to its status as one of the few results in perception science that has captured public interest and imagination (Chabris & Simons, 2011; Cloud, 2010; Murphy, 2017).

Crucially, however, this interpretation of IB and the many implications that follow from it rest on a measure that psychophysics has long recognized to be problematic: simply asking participants whether they noticed anything unusual. In IB studies, awareness of the unexpected stimulus (the novel shape, the parading gorilla, etc.) is retroactively probed with a yes/no question, standardly, “Did you notice anything unusual on the last trial which wasn’t there on previous trials?”. Any subject who answers “no” is assumed not to have any awareness of the unexpected stimulus.

However, yes/no questions of this sort are inherently and notoriously subject to bias, because they require observers to set a *criterion* in order to decide whether they have enough evidence to answer “yes” or instead answer “no” (Dulany, 2001; cf. Eriksen, 1960; Holender, 1986; Irvine, 2012). Under the framework of Signal Detection Theory (Tanner & Swets, 1954; Green & Swets, 1966), observers who are asked to determine whether a signal is or is not present must in setting such a criterion consider tradeoffs between the various possible outcomes of their decision (i.e., the relative costs of hits, misses, false alarms, and correct rejections). Indeed, in an IB task (where the signal in question is the unexpected stimulus), observers may have reason to be conservative in their criterion-setting—i.e., to adopt a high standard for saying “yes” (and answer “no” if that standard isn’t met). For example, observers might be under-confident whether they saw anything (or whether what they saw counted as unusual); this might lead them to respond “no” out of an excess of caution. Subjects might doubt that they could identify it if asked, and so respond “no” to avoid having to do so. Subjects may also worry that if they report noticing the unexpected object, the experimenters will take that to mean they weren’t engaged in the task they had been given (e.g., judging which cross arm was longer or counting the passes; Dulany, 2001). On any of these possibilities, subjects who say they did not notice the critical stimulus may well have had some awareness of it, but simply underreported it given the constraints of traditional IB questioning.

Evidence for this alternative hypothesis would have dramatic consequences for the dominant interpretation of IB and the implications that follow from it: It would overturn the view that IB reflects a total failure to perceive and so challenge the appeal to IB in arguments that attention is required for awareness. Remarkably, however, the hypothesis that subjects in inattentional blindness tasks actually see more than they say (but respond otherwise because they are conservative in reporting their awareness) remains empirically unsettled, and the powerful tools of signal detection theory unexploited in relation to IB.

A handful of prior studies have explored the possibility that inattentionally blind subjects may retain some visual sensitivity to features of IB stimuli (e.g., Schnuerch et al., 2016; see also Kreitz et al., 2020, Nobre et al., 2020). However, a recent meta-analysis of this literature (Nobre et al., 2022) argues that such work is problematic along a number of dimensions, including underpowered samples and evidence of publication bias that, when corrected for, eliminates effects revealed by earlier approaches, concluding “that more evidence, particularly from well-powered pre-registered experiments, is needed before solid conclusions can be drawn regarding implicit processing during inattentional blindness” (Nobre et al., 2022).

Here, five experiments provide this critical test for the first time. We achieve this by making four key modifications to classic IB paradigms. First, we add follow-up questions to the classic yes/no methodology to explicitly probe awareness of features of the IB stimulus (i.e., not just asking whether subjects noticed anything, but also querying features of the object they said they didn’t notice). Although the use of follow-up questions is not new, our approach overcomes several problems with their past use in the literature (see General Discussion for much more on this point). Second, we include “absent trials” in which some subjects are shown no additional stimulus on the critical trial but are still asked whether they noticed anything unusual. This procedure provides a false alarm rate to test—for the first time, to our knowledge—whether subjects are conservative in reporting their awareness in the ways hypothesized above. Third, we leverage online data collection to massively increase sample size (running 25,000 subjects, each experiencing just a single critical trial), in order to make possible the crucial signal-detection analyses that separate bias from sensitivity. Finally, to overcome the fact that in IB each subject necessarily only contributes one trial (since following a critical trial, stimuli are no longer unexpected), we introduce a novel analytic approach by applying signal detection models to a “super subject”.

Altogether this approach reveals that as a group subjects can report above-chance the features of stimuli (color, shape, and location) that they had all claimed not to notice under traditional yes/no questioning, and that this underreporting of awareness is well-modeled in terms of a conservative criterion. In place of the dominant interpretation that IB abolishes perception and awareness, the present results indicate that significant residual sensitivity remains in IB, and are even consistent with an alternative picture on which inattention instead *degrades* awareness. More generally, our findings motivate an approach to perception and awareness which treats them as graded as opposed to all-or-nothing, which we further discuss below.

## Results

### Above-chance sensitivity to inattentional blindness stimuli: Location

Experiment 1 modified the canonical inattentional blindness task used by Mack and Rock (1998; Figure 2a). On three trials, subjects were presented with a cross randomly assigned on each trial to be directly above or below a central fixation point (for 200 msec), and their task was to report which arm of the cross (horizontal or vertical) was longer. The fourth, critical trial, proceeded in the same way, but with the addition of an unexpected red line appearing in the periphery simultaneous with the cross. After again reporting which arm of the cross was longer, subjects were asked the standard question used to measure inattentional blindness: “Did you notice anything unusual on the last trial that wasn’t there on previous trials?” (yes or no). In line with established findings on IB, a considerable proportion of subjects (28.6%) responded “no” they didn’t notice anything unusual (we refer to these subjects across all our experiments as ‘non-noticers’).

**Figure 2.**
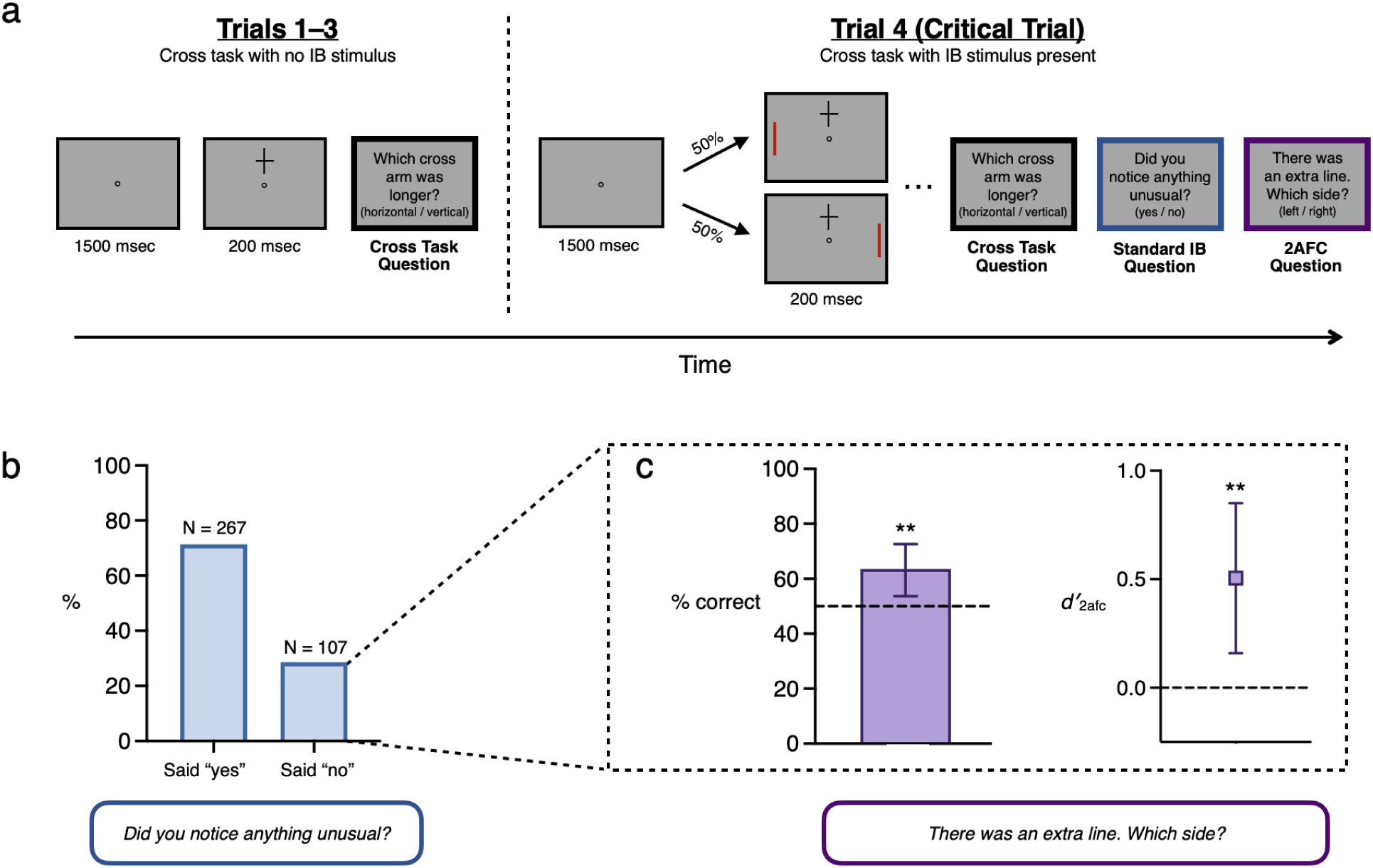
Stimuli, procedure and results for Experiment 1. **a** Schematic trial sequence for Experiment 1. On trials 1-3 subjects were presented with a cross above or below fixation for 200 msec and judged which arm was longer. On trial 4, an unexpected red line appeared in the periphery simultaneous with the cross. After reporting which cross-arm was longer, subjects were asked, “Did you notice anything unusual on the last trial that wasn’t there on previous trials?” (yes/no), followed by a 2afc question concerning the location of the line (left/right). **b** Percentage of subjects who report noticing or not noticing the extra red line. 29% of subjects were ‘non-noticers’. **c** Performance on the 2afc question (left/right location of the line), considering only those subjects who reported not noticing anything unusual. Remarkably, both %-correct and *d′* were significantly above chance among the group of subjects who met traditional criteria for inattentional blindness. In other words, collectively, subjects who answered “no” demonstrated sensitivity to the location of the stimulus they all had just claimed not to have noticed. Error bars are 95% CIs.

Following the standard IB question, our modification included an additional question: “In fact, the last trial you just saw contained one extra element—a vertical red line on either the left or the right side of the box. What side do you think it appeared on?” (left or right). Far more subjects were able to correctly locate the stimulus (87.4%) than said they noticed it (71.4%), raising the possibility that non-noticers—i.e., those who demonstrated inattentional blindness—might have performed significantly better than chance. This is precisely what we found: 63.6% of non-noticers answered the forced-choice question correctly (significantly different from the 50% correct expected by chance in a two-sided binomial probability test, 95% CI = [0.54, 0.73], BF10 = 9.9). In other words, nearly two-thirds of subjects who had just claimed not to have noticed any additional stimulus were then able to correctly report its location.

Critically, this result also holds using *d′*, an unbiased measure of sensitivity (*d′_2afc_* = 0.51, 95% CI = [0.16, 0.85]; see Box 1 on Signal Detection Theory), which we use in reporting all following results concerning sensitivity in non-noticers. An important novelty of our strategy is that it derives these statistics in relation to a “super subject” whose responses are comprised of individual subjects’ responses in their single critical trials. Note that all analyses reported here relate to this super subject as opposed to individual subjects; see General Discussion for more on the assumptions behind this analytic approach.

#### Box 1: Signal Detection Theory and Inattentional Blindness

In Signal Detection Theory (SDT; Green and Swets, 1966; Macmillan and Creelman, 2005), perceptual decisions are based on statistically variable sensory evidence (*S + N*) due to a stimulus or signal (*S*), if any, together with omnipresent noise (*N*). Deciding whether a stimulus was present or not (yes/no detection) involves deciding whether the evidence received is a sample from *S + N* or from *N* alone. To make this decision, an observer must set a *criterion*, a level of sensory evidence sufficient for a positive response. This criterion is flexible and can be adjusted in accord with the payoffs associated with different outcomes such as hits (saying “yes” when a stimulus is present) and false alarms (saying “yes” when no stimulus is present).

SDT uses observed hit and false alarm rates to distinguish two distinct aspects of an observer’s performance: their *sensitivity* and *bias*. In simple models, *N* and *S + N* distributions are treated as equal variance Gaussians. The *sensitivity* of an observer is then measured as the standardized distance between the means of the *N* and *S + N* distributions. Intuitively, if the two distributions entirely overlap, an observer is entirely insensitive to the presence of a stimulus, whereas the greater the separation, the greater the sensitivity. To calculate this measure of sensitivity, we subtract the *z*-transform of the false alarm rate from the *z*-transform of the hit rate: *d’ = z*(*H*) - *z*(*FA*). Critically, this measure is independent of the location of a subject’s criterion and so offers an objective, bias-free measure of their sensitivity.

The *bias* of an observer is their tendency to prefer a particular response independent of the actual presence of a stimulus. This can be measured by the location of their criterion with respect to the midpoint of the *N* and *S + N* distributions. Intuitively, a criterion at the midpoint shows no preference for “yes” responses over “no” responses independent of the actual presence of a stimulus, whereas a *conservative* criterion reflects a preference for “no” responses, and a *liberal* criterion a preference for “yes” responses. We can calculate a standardized measure of bias, as follows: *c =* -½[*z*(*H*) + *z*(*FA*)].

The traditional question in IB studies—“Did you notice anything unusual on the last trial that wasn’t there on previous trials?”—can be treated as a yes/no detection question. By including absent trials, where no stimulus was presented and yet we ask this question anyway, we determined hit and false alarm rates across critical trials after applying the Hautus (1995) correction of adding 0.5 to every cell in the decision matrix to avoid infinite values which would arise if any cell were zero. For example, in our Exp. 5, 7.2% of subjects who were shown no IB stimulus nonetheless responded “yes”—yielding a corrected false alarm rate of 0.073. On the other hand, 71% of subjects who were shown a stimulus responded “yes”—yielding a corrected hit rate of 0.71. Using the calculations described, we could separate the sensitivity of the corresponding super-subject from their bias. This revealed a significant *conservative* bias (*c =* 0.45), indicating a preference to deny noticing an unexpected stimulus, independent of its actual presence.

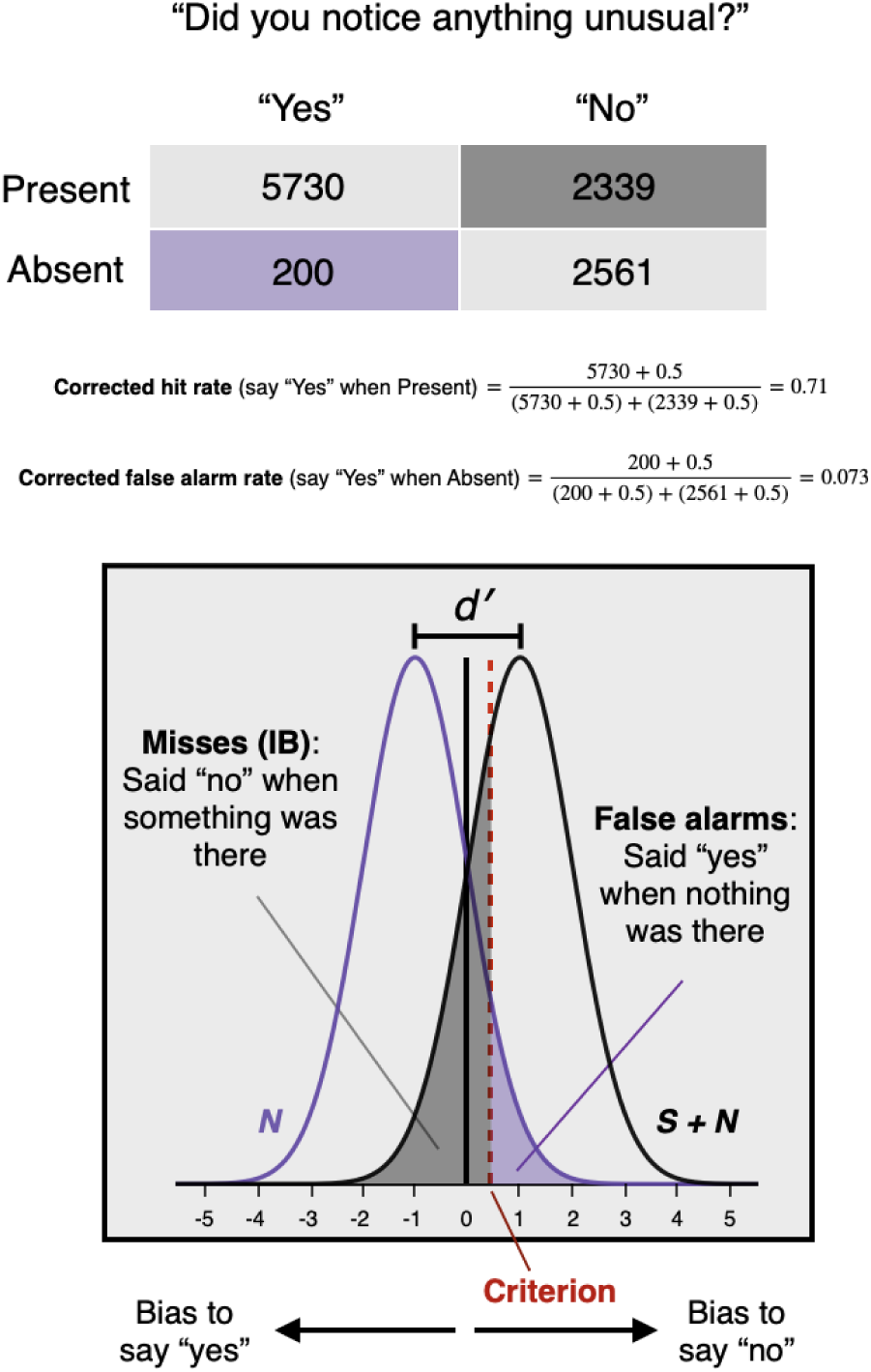

### Above-chance sensitivity to inattentional blindness stimuli: Color

Experiment 2 repeated the design of Experiment 1, except that the additional line could be either red or blue, and the question about the line’s location was replaced with a one-interval forced-response question about the line’s color. This experiment yielded similar results to Experiment 1: Only 72.3% of subjects shown an additional stimulus said they had noticed something, yet 81% were able to indicate the additional line’s color correctly. And again, as a group, non-noticers demonstrated above chance sensitivity, this time to color (*d′* = 0.38, 95% CI = [0.03, 0.73])^1^—note that the unbiased nature of this measure is especially critical here since subjects displayed a significant bias in favor of responding “blue” (*c* = 0.67, 95% CI = [0.49, 0.84]). In other words, consistent with the results of Experiment 1, as a group, subjects who had just claimed not to have noticed an additional stimulus were able to correctly report its color at rates well above chance. This pair of initial results shows that even subjects who answer “no” under traditional questioning can, as a group, still correctly report various features of the stimuli they just reported not having noticed, indicating significant group-level sensitivity to visual features. Moreover, these results are even consistent with an alternative hypothesis about IB, that as a group, subjects who would traditionally be classified as inattentionally blind are in fact at least partially aware of the stimuli they deny noticing.

### Conservative reporting of visual awareness

Our results raise a natural question: Why, if subjects could succeed at our forced-response questions as a group, did they all individually claim not to have noticed anything? Experiment 2 made an additional modification precisely to address this question: the introduction of ‘absent’ trials in which no additional line was shown but subjects were still asked the yes/no and one-interval forced-response questions. Absent trials provide an additional source of information about subjects’ biases, by revealing how often they respond “yes” without any stimulus present (i.e., their false alarm rate). This allows for the computation of response bias (*c*) in relation to the crucial IB question(“Did you notice anything unusual on the last trial that wasn’t there on previous trials?” again, see Box 1), which to our knowledge is unique in this literature. This analysis revealed evidence for the second aspect of our alternative hypothesis: Not only do subjects collectively have residual sensitivity to unnoticed IB stimuli, they are, as a group, *conservative* in reporting their awareness (*c* = 0.31, 95% CI = [0.22, 0.41]; *d′* = 1.81, 95% CI = [1.63, 1.99]; note that statistics here are for all subjects). In turn, this raises the possibility that subjects may retain *awareness* of unreported IB stimuli corresponding to their residual visual sensitivity but that this awareness is systematically underreported.

### Above-chance sensitivity in high-confidence non-noticers

Although the previous two studies are consistent with the hypothesis that “inattentionally blind” subjects retain at least partial awareness of unattended stimuli (and are conservative in reporting that awareness), it is possible that these results were driven by a subset of subjects, with other subjects remaining truly blind to the IB stimulus (i.e., having no sensitivity at all to its features). This might arise if above-chance sensitivity were restricted to subjects who were under-confident in responding “no” when asked whether they had noticed anything unusual. In that case, subjects who *confidently* answered “no” (i.e., felt certain they didn’t notice any additional stimulus) might, as a group, fail to perform above chance on the subsequent discrimination task. Experiment 3 addressed this possibility directly, by (a) adding confidence ratings to the standard yes/no question, and (b) dramatically increasing the sample size, so as to separately analyze the performance of high- and low-confidence subjects. The task proceeded in the same way as Experiment 1, except that after the yes/no question about noticing anything unusual, subjects were asked to rate their confidence in their answer, on a 4-point scale from 0–3 (0 = Not at all confident; 3 = Highly confident); finally, subjects were then asked the left/right discrimination question (and gave their confidence in that answer as well, though this was less crucial to our hypothesis—see Supplementary Materials for details). As shown in Figure 3e, answers to these questions ran the full spectrum of responses, with subjects expressing varying degrees of confidence in “yes” and “no” responses to the IB question. Of particular relevance was the group of “high-confidence non-noticers”—i.e., subjects who said “no” (they didn’t notice anything unusual), and then rated their confidence in that answer as “3” (highly confident). Remarkably, even this group of subjects (N = 204) collectively demonstrated significantly above-chance sensitivity to the location of the IB stimulus (*d′_2afc_* = 0.34; 95% CI = [0.08, 0.60]). Further, as is evident from Figure 3e, subjects’ confidence in their yes/no response predicted accuracy for that group on the discrimination task, suggesting that subjects may have graded awareness of unattended stimuli in IB tasks (see Supplementary Materials for details).

**Figure 3.**
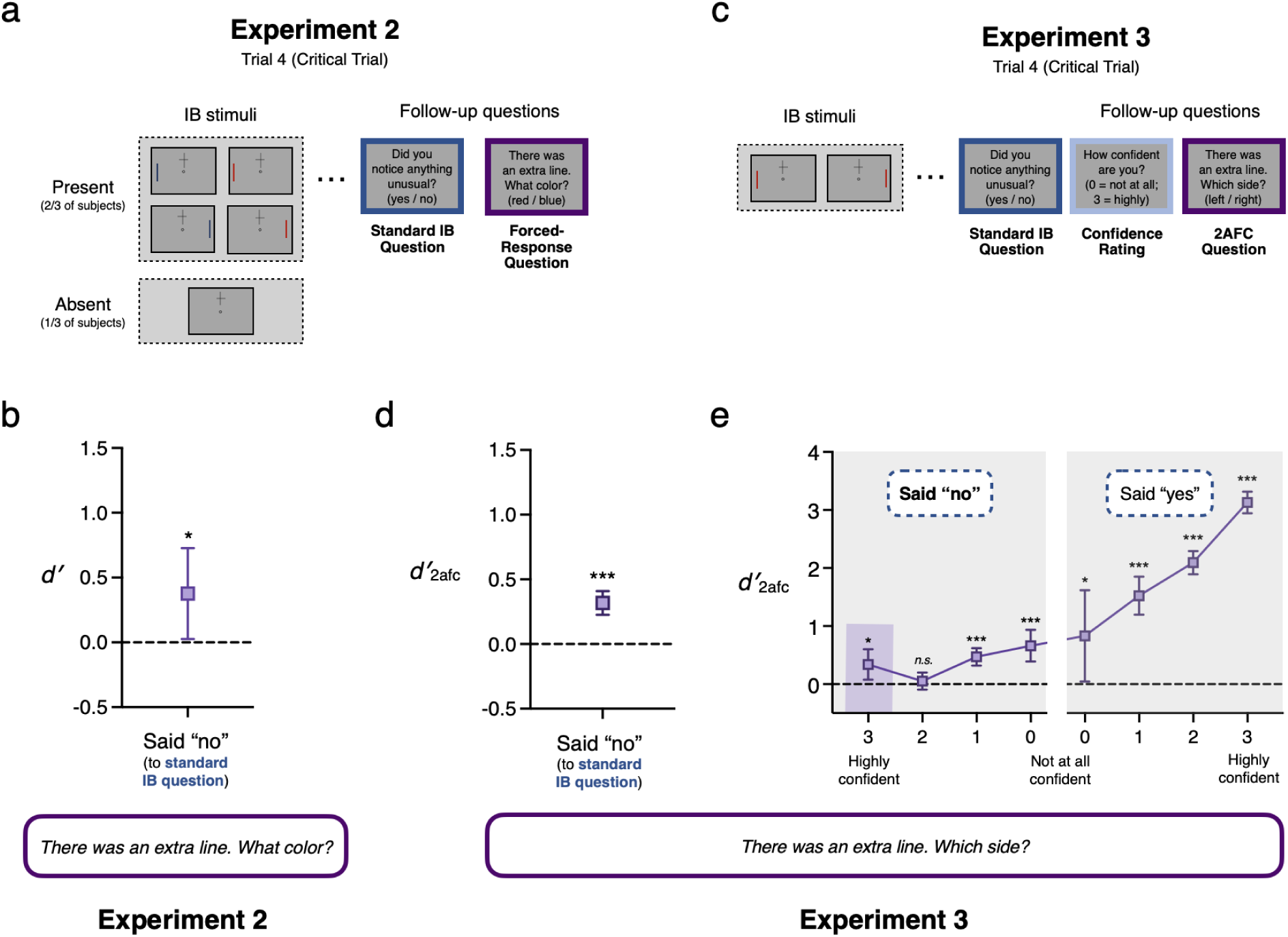
Stimuli, procedure and results for Experiments 2 and 3. **a** Schematic trial sequence for Experiment 2. On trials 1-3 subjects were presented with a cross above or below fixation for 200 msec and judged which arm was longer. On trial 4, for 2/3rds of subjects, an unexpected line appeared in the periphery simultaneous with the cross. This line could be either blue or red and one the left or right. For 1/3 of subjects no additional line was shown. After reporting which cross-arm was longer, all subjects were asked, “Did you notice anything unusual on the last trial that wasn’t there on previous trials?” (yes/no), followed by a one-interval forced-response question concerning the color of the line (red/blue). **b** Performance on the one-interval forced-response question about the unexpected line’s color amongst subjects who were shown a line and who reported not noticing anything unusual (27.73% of subjects). As a group, subjects who answered “no” demonstrated sensitivity to the color of the stimulus they had just claimed not to have noticed. **c** Schematic trial sequence for Experiment 3. Trials 1-4 were identical to Experiment 1, except subjects were asked additional questions about their confidence following both yes/no and 2afc questions. **d** Performance on the 2afc question in Experiment 3, considering only subjects who reported not noticing anything unusual (30.85% of subjects). Replicating the finding of Experiment 1, as a group, subjects who answered “no” demonstrated sensitivity to the location of the stimulus they had just claimed not to have noticed. **e** Performance on the 2afc question in Experiment 3 for all subjects, broken down by confidence in their response to the yes/no question whether they had noticed anything unusual. Remarkably, even subjects who were highly confident that they had not noticed anything unusual were collectively significantly above chance. Error bars are 95% CIs.

### Generalizing to dynamic inattentional blindness

Experiments 1-3 suggest that, collectively, subjects underreport their perception of a brief (200 msec) IB stimulus. However, in classic studies of dynamic inattentional blindness (Simons & Chabris, 1999; Most et al., 2005; Ward & Scholl, 2015), the unexpected stimulus remains in view for an extended period. Here, it is tempting to think that subjects will not be conservative in reporting that they noticed an IB stimulus given they have many seconds to build confidence in what they saw. Experiment 4 tested this possibility by modifying a traditional sustained inattentional blindness paradigm in which the IB stimulus remains on screen for 5 seconds.

Finding the same pattern of above-chance sensitivity and conservative response bias in this very different paradigm would be striking evidence that, quite generally, there is residual sensitivity to visual features in inattentional blindness, and lend further support to our alternative hypothesis.

In Experiment 4, subjects’ primary task was to count how often squares of a particular color (black or white) bounced off the perimeter of a gray rectangle (adapted from Wood & Simons, 2017, in turn based on Most et al., 2001; see Figure 4a). This task demands significant attention as each set of colored squares bounces an average of 28 times during a 17-second trial. For some subjects, on the third and critical trial, an additional shape (a triangle or circle, which was either black or white) entered the display (on the left or the right) and traversed the full height of the display in a straight path (from either top to bottom, or bottom to top), remaining on screen for 5 seconds before disappearing. When the trial ended, subjects reported how many times the squares of their assigned color bounced. They were then immediately asked the standard question used to measure inattentional blindness: “Did you notice anything unusual on the last trial that wasn’t there on previous trials?” (yes or no). Following this, each subject answered *three* additional questions about the extra object that may or may not have appeared, in a random order: (1) What color was it? (black or white), (2) What shape was it? (circle or triangle), (3) What side was it on? (left or right).

**Figure 4.**
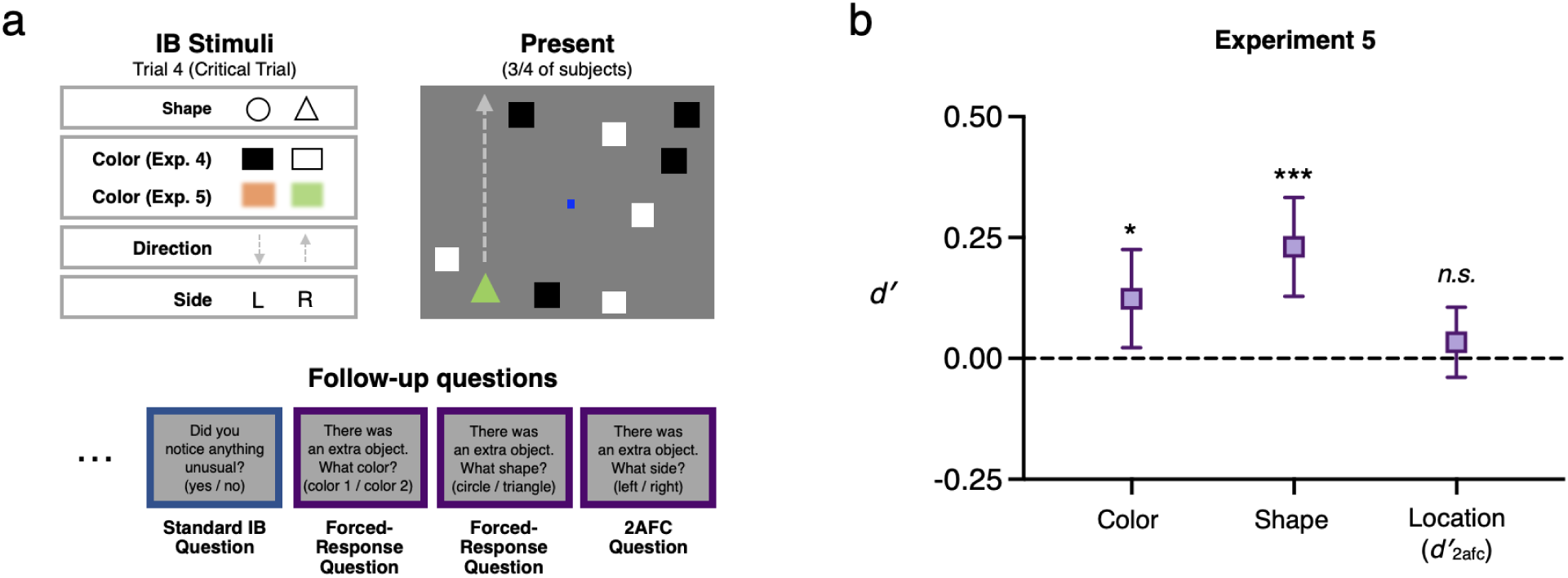
Stimuli, procedure and results for Experiment 5. **a** Stimulus parameters and a schematic critical trial for Experiment 5, in which an unexpected object moved across the display for several seconds while subjects counted the number of times the white squares bounced off the display’s ‘walls’ (task adapted from Wood & Simons, 2017). The unexpected object varied in its shape, color, direction of motion, and side of the display. As in Experiment 4, subjects were then asked follow-questions about this stimulus’s color, shape and location. **b** As a group, subjects who reported not noticing the unexpected stimulus still showed above-chance sensitivity to its color and shape (though not to its location), a pattern predicted in our pre-registration. Thus, sensitivity to IB stimuli arises even when the stimuli are visible for a sustained period (rather than appearing only briefly, as in Experiment 1-3). Error bars are 95% CIs.

**Figure 5.**
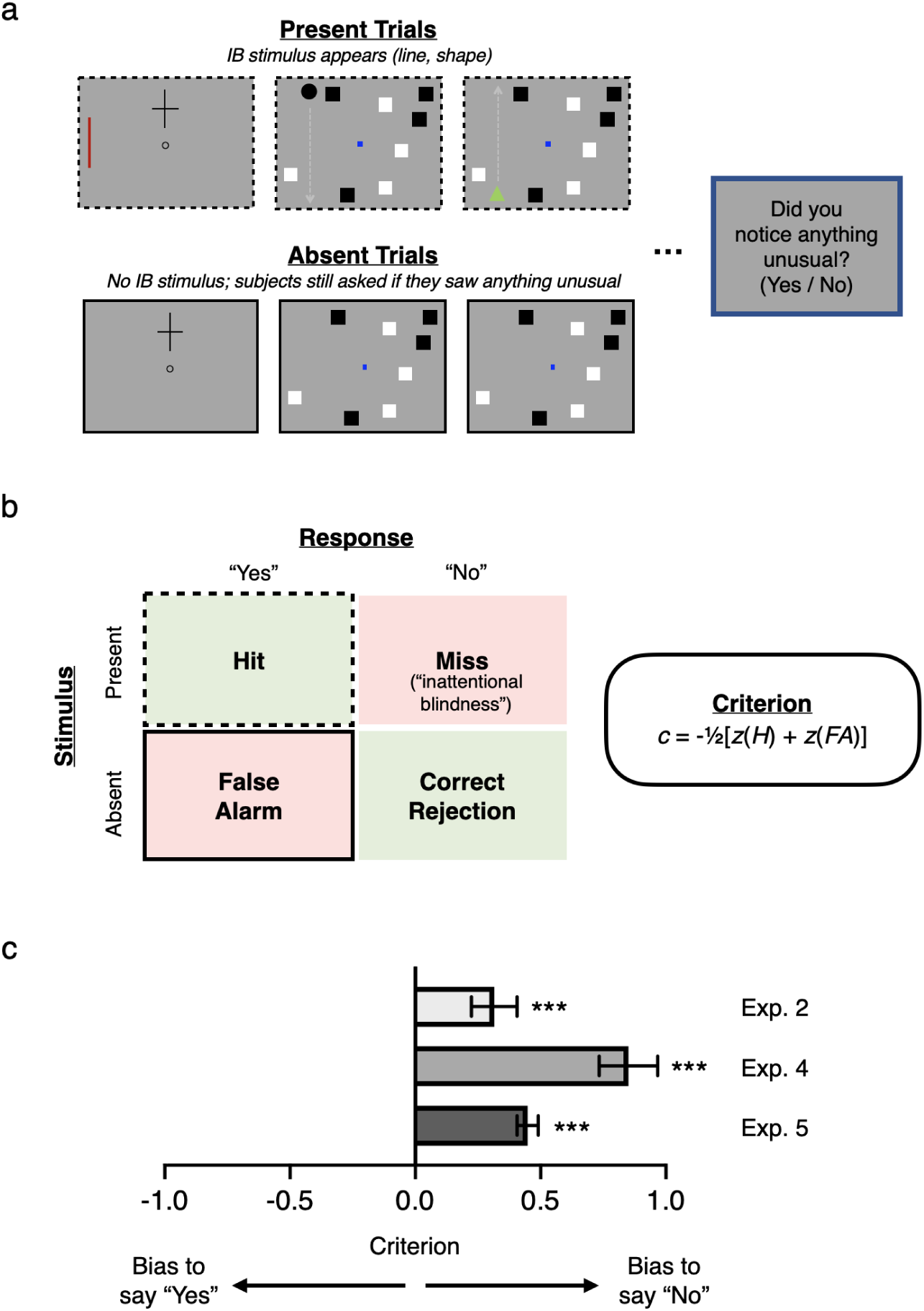
Conservative criteria in Experiments 2, 4 and 5. **a** Critical trials from Experiments 2, 4 and 5, showing present trials in which an IB stimulus was presented to subjects (2/3rds subjects in Experiment 2, 3/4 in Experiments 4 and 5), and absent trials in which no IB stimulus was presented (1/3rd subjects in Experiment 2, 1/4 in Experiments 4 and 5). On all trials, subjects were asked, “Did you notice anything unusual on the last trial that wasn’t there on previous trials?” (yes/no). **b** Left: Decision matrix for this yes/no question, indicating the four possible stimulus/response pairings: hits corresponding to ‘yes’ responses on present trials; false alarms corresponding to ‘yes’ responses on absent trials; misses corresponding to ‘no’ responses on present trials (i.e. inattentional blindness); and correct rejections corresponding to ‘no’ responses on absent trials. Right: Formula for calculating criterion or response bias from hit rate (H), i.e. proportion of present trials in which subjects responded ‘yes’, and false alarm rate (FA), i.e. proportion of absent trials in which subjects responded ‘no’. **c** Criteria for Experiments 2, 4 and 5, showing that in each case subjects exhibited a significantly conservative bias, i.e. a tendency to say ‘no’ when asked if they noticed anything unusual, independent of the actual presence of a stimulus. This suggests that, collectively, subjects in inattentional blindness experiments may systematically underreport their awareness of unexpected stimuli across different paradigms.

Consistent with previous studies using dynamic stimuli (see Figure 1b and 1c), a majority of subjects—57.3%—demonstrated IB. Put another way, just 42.7% of subjects who were shown something additional on the critical trial were correct on the yes/no detection question. However, despite the attentionally demanding primary task, non-noticers collectively again demonstrated significant sensitivity to the features of the IB stimulus, choosing the correct color (*d′* = 0.82, 95% CI = [0.61, 1.04]) and shape (*d′* = 0.21, 95% CI = [0.01, 0.42]) significantly more often than would be predicted by chance. Given that all the objects were changing location constantly in Experiment 4, we did not pre-register a prediction about location discrimination being above-chance, and we did not find that non-noticers performed above chance on the left/right discrimination question (though we did observe a trend in this direction, *d′*_2afc_ = 0.07, 95% CI = [-0.08, 0.22]).

Just as in Experiment 2, subjects underreported their awareness of the IB stimulus on the traditional yes/no question, employing a decision criterion that was even more conservative than that measured in Experiment 2 (*c* = 0.85, 95% CI = [0.73, 0.97]; *d′* = 1.33, 95% CI = [1.10, 1.57]; again, note that statistics here are for all subjects). The fact that subjects were collectively more reluctant to report their awareness of the IB stimulus in this experiment compared with Experiment 2 is surprising given that subjects had 5 full seconds to build confidence that there was a stimulus. However, given that the colors of the squares in the primary task were the same as the colors used for the IB stimuli, it is possible that estimates of criterion and color sensitivity in Experiment 4 were affected by an interaction effect due to enhancement of color-congruent stimuli and suppression of color-incongruent stimuli (cf. Wood & Simons, 2017). Specifically, the IB rate for color-incongruent stimuli was substantially higher than for color-congruent stimuli (86.57% vs. 28.71%), suggesting suppression occurred. That is, a subject shown an unexpected black stimulus while attending to black squares was more likely to report noticing than a subject shown an unexpected white stimulus. This in turn may have inflated our estimate of color sensitivity if subjects correctly assumed that they would have been more likely to notice a color-congruent stimulus and biased their responses towards the incongruent color.

To address this, Experiment 5 repeated the design of Experiment 4, with two key differences: (1) Every subject was assigned to attend to the white squares in the primary task, (2) the IB stimuli were either orange or green—two colors that as a pair produced equal IB rates in pilot testing. Moreover, by selecting such highly salient colors (which differ dramatically from any other stimulus on the display), these parameters served as an especially strong test of our broader hypothesis. Remarkably, even under these conditions (and with the concern about color congruency effects mitigated), the pattern of results matched that of Experiment 4: Subjects collectively performed above-chance in discriminations of both color (*d′* = 0.12, 95% CI = [0.02, 0.23]) and shape (*d′* = 0.23, 95% CI = [0.13, 0.33]), but not location (*d′_2afc_* = 0.03, 95% CI = [-0.04, 0.11]). Sensitivity to the color of the IB stimulus was lower than in Experiment 4, consistent with our prediction that color sensitivity in that experiment was inflated by a color-congruence bias; however, it was still significantly above chance, indicating that color-congruence does not fully account for the pattern of results. That subjects,as a group, could still correctly report the color and shape of the IB stimulus despite the attentionally demanding primary task is compelling evidence that perception of IB stimuli is not completely abolished by inattention. Further, we again found that subjects were collectively biased to respond “no” to the traditional yes/no IB question (*c* = 0.45, 95% CI = [0.41, 0.49]; *d′* = 2.01, 95% CI = [1.94, 2.09]; statistics here are for all subjects). Thus, even when an unexpected stimulus remains on screen for many seconds, subjects are hesitant to report noticing anything unusual.

## General Discussion

Inattentional blindness captures both scholarly interest and popular imagination because of its striking and counterintuitive implication that we may fail to see what is right before our eyes, simply because our attention is otherwise engaged. Its influence is both wide and deep: It apparently provides a dramatic demonstration of the limits of visual perception, serves as a tool to reveal the neural correlates of consciousness, and even motivates theories of consciousness holding that awareness requires attention. These and still other implications arise from the “consensus” interpretation of IB, according to which inattention completely abolishes perception of the unexpected stimulus: “one can have one’s eyes focused on an object or event … without seeing it *at all*” (Carruthers, 2015, emphasis added). Yet this interpretation rests on a crucial and untested assumption: that observers who say they didn’t notice a stimulus (i.e., answer “no” under traditional yes/no questioning) in fact didn’t see it.

The present work puts this crucial assumption to the test, yielding results that point to a very different pattern than the consensus interpretation: Across 5 pre-registered experiments totaling over 25,000 subjects, we found that groups of observers could successfully report a variety of features of unattended stimuli, even when they all individually claimed not to have noticed those stimuli. Furthermore, our approach revealed that subjects are conservatively biased in reporting their awareness, in ways that not only explain our results (i.e., provide an account of how and why subjects who claimed not to have seen something could still report its features) but also recast the large and influential body of literature that has taken answers to yes/no questions in IB paradigms at face value.

### Design and analytical approach

Our experiments all employed designs and protocols closely modeled on canonical IB studies. In Experiments 1-3, we studied IB using a cross task closely modeled on Mack and Rock’s classic studies (Mack & Rock, 1998). In Experiments 4-5, we studied IB using a dynamic task closely modeled on Wood & Simons (2017) (itself adapted from influential work by Most et al., 2001; Most et al., 2005; Ward & Scholl, 2015). In all cases, we used the standard yes/no question from previous experiments to determine IB rates. These choices were deliberate: Our aim was to interrogate the canonical interpretation of a large and long-standing tradition of experimental work, and so we sought to cleave as closely as possible to the experiments which inspired and have been subject to this interpretation. Our results do not reflect idiosyncratic design choices but rather speak to the central paradigms in the literature.

Our approach also eschews the use of divided and full attention trials. Divided attention trials (trials after the critical trial in which the subject knows that an unexpected stimulus might occur) and full attention trials (where in addition the subject is told to ignore the primary task, and instead look out for unexpected objects) are often used to exclude from analysis subjects who do not report seeing the unexpected stimulus on these trials (see, e.g. Most et al. 2001). However, this practice is controversial and has been argued to “lead researchers to understate the pervasiveness of inattentional blindness” (White et al., 2018). Because our aim was to offer an especially stringent test of sensitivity in inattentional blindness, we opted not to use any such exclusions. Since the use of such exclusions would tend to inflate the biases we are concerned with here, the fact that we found evidence of residual sensitivity despite not excluding subjects in this way is all the more telling.

To assess objective sensitivity and bias in IB, we adopted a novel analytical approach, applying signal detection models to a “super subject” whose responses are comprised of individual subjects’ responses in their single critical trials. This strategy involves various assumptions which future work should explore. Nonetheless, we believe any violations of such assumptions will not affect our main results or, where they might, lead only to our underestimating residual sensitivity and response bias. First, in calculating *d′*, it is standardly assumed that a subject’s criterion is stable throughout a given experiment. This assumption may be more likely to be violated with respect to a “super subject”. However, its violation will not affect calculations of *d′* in 2afc tasks which are criterion free (Exps. 1, 3 and location results in Exps. 4 & 5). Moreover, if criterion instability is present, its effect will be to reduce estimated sensitivity in one-interval forced-response tasks (Exp. 2, and shape and color results in Exps. 4 & 5; see, Azzopardi & Cowey, 2001). Our approach can thus be seen as offering a conservative estimate of residual sensitivity. (Indeed, given the essentially retrospective nature of IB judgments, our estimates should in any case be considered conservative since signal available at the time of stimulus presentation may have been lost by the time of judgment.) Second, in calculating *d′* and *c*, it is standardly assumed that signal and noise distributions are equal variance Gaussians. There is theoretical reason to think this assumption is robust with respect to 2afc tasks (Macmillan & Creelman, 2005), and in general in relation to the Gaussian nature of the distributions (Green & Swets, 1966; Pastore et al., 2003; Wixted, 2020). However, empirically, equal variance might not hold in one-interval tasks. Violation of this equal variance assumption could lead to under- or over-estimation of *d′* and *c.* However, any such under- or over-estimation would be slight and unlikely to affect our main results. For example, if the relative variance of the signal-plus-noise distribution as compared to the noise distribution (*σ*) is 1.25, then *c* for the yes/no task in Experiment 5 (illustrated in Box 1) would be 0.443 and *d′* would be 1.898; whereas, if *σ* = 0.75, then *c* would instead be 0.438 and *d′* would be 2.118. In either case, our main results are qualitatively unchanged.

Our super subject analysis raises the question of how to interpret the responses of individual subjects. Even though our experiments revealed that subjects who denied noticing any unusual stimulus could *collectively* report its features above chance, it was also the case that some subjects denied noticing any unusual stimulus and then also went on to incorrectly answer the follow-up questions about its features. Indeed, this has also been true in other studies that include follow-ups (e.g., Most et al., 2001, 2005; Cohen et al., 2011, 2020). Are these individual subjects truly inattentionally blind? Intriguingly, even this seemingly cautious conclusion does not follow from that pattern of performance, for several reasons. First, and straightforwardly, many such follow-ups are not 2afc questions but rather yes/no questions themselves (including our own questions about color and shape, as well as many similar questions used in previous work); since such questions are biased (e.g., subjects may tend to favor responding that an object was blue rather than red, perhaps making assumptions about the visibility of each color), any individual incorrect answer to such a question may reflect this sort of bias as opposed to the total absence of color signal. Second, even in an unbiased 2afc task, an observer may have significant information from the stimulus (perhaps, well above an unbiased single-interval detection criterion) but still decide incorrectly because of high noise from the other spatial interval; in such a case, it is far from clear that the subject should be treated as blind. Third, there will inevitably be subjects who fail to correctly report an unnoticed object’s features because they failed to see it in the first place — not due to inattention, but rather due to more ordinary failures such as happening to look away from the display at the key moment, sneezing or blinking just as the unexpected stimulus appears, being interrupted by one’s child or pet or smoke alarm, and so on; such subjects would be “blind” to the stimulus, of course, but not *inattentionally* blind. Fourth any series of follow-up questions, including ours, inevitably probes only some limited set of features at the exclusion of others; thus, subjects may have been aware of some feature of the stimulus other than the features explicitly probed (e.g., the orientation of an unexpected line rather than its color). Fifth, many processes intervene between being (or not being) sensitive to a stimulus and generating a response to a follow-up question; subjects will occasionally press the wrong button by mistake, or rush through the questions without reading carefully, or forget what they saw (and thus guess); though such mishaps will tend to break in favor of incorrect answers just as often as correct answers, they make it so that any individual error (or success) is difficult to interpret on its own (and in ways that testify to the value of the group-level approach that we favor here). Sixth, and perhaps most generally, taking correct and incorrect answers to place subjects neatly into two categories — those who saw the stimulus and those who did not — reflects a binary approach to perception and awareness that we suggest should be resisted. Indeed, an aspect of our contribution here, discussed further below, is to encourage conceiving perception and awareness as coming in degrees, in line with a Signal Detection Theory framework. On this view, perception and awareness are most helpfully characterized in terms of continuous statistics such as *d′* rather than more traditional but problematic measures such as the proportion of correct or incorrect responses.

### Relation to previous work

Our work was motivated by concerns that the traditional interpretation of IB relies on assessing perception and awareness simply by asking participants whether they noticed anything unusual. As discussed, such yes/no questions are notoriously subject to bias, which may lead subjects to answer “no” even when they do have a degree of perception or awareness, due to factors such as underconfidence. More recent studies of IB have attempted to improve on simple yes/no questioning through the use of various follow-up questions. However, although these improved methods undoubtedly have their merits, none of them resolves the concerns that motivated our investigations. This is for three fundamental reasons.

First, follow up questions are often used not to *exclude* subjects from the IB group but to *include* subjects. For example, Most et al. 2001 treated as inattentionally blind not only subjects who denied noticing the unexpected object but also subjects who claimed they did notice the object but were subsequently unable to describe it. Similarly, Pitts et al. (2012) asked subjects to rate their confidence in their initial yes/no response from 1 = least confident to 5 = most confident, and used these ratings to include in the IB group those who rated their confidence in seeing at 3 or less. Counting such subjects as inattentionally blind can be problematic. There is a large gap between being under confident that one saw something and being completely blind to it; failure to describe a feature (e.g., color, shape) does not imply a complete lack of information concerning that feature; and even if a subject did lack all information concerning some feature of an object, this would not imply a complete failure to see the object.

Second, most follow up questions remain subject to response bias in just the same way as the original yes/no awareness question. For example, Cohen et al. (2020; see similarly: Cohen et al., 2011; Simons et al., 1999; Most et al., 2001, 2005; Drew et al., 2013; Memmert, 2014) use a series of follow up probes: (1) “Did you notice anything strange or different about that last trial?” (2) “If I were to tell you that we did something odd on the last trial, would you have a guess as to what we did?” (3) “If I were to tell you we did something different in the second half of the last trial, would you have a guess as to what we did?” (4) “Did you notice anything different about the colors in the last scene?” These questions are intrinsically subject to response bias just by being yes/no questions; but in this case they may be especially susceptible since subjects may be reluctant to ‘take back’ their earlier answers, leaving them all the more conservative to avoid any perceived inconsistency. (This may also explain the remarkable consistency in such responses reported in, e.g., Simons & Chabris, 1999, despite the very different wording across the questions asked.) It is also important to recognize that whereas 2afc questions are criterion free (in that they naturally have an unbiased decision rule), this is not generally true of *n*afc nor delayed *n*-alternative match to sample designs. Performance in such tasks thus requires SDT analysis – which itself may be problematic if the decision space is not properly understood or requires making substantial assumptions about observer strategy.

Third, and finally, many follow up questions are insufficiently sensitive (especially with small sample sizes). For instance, Todd, Fougnie & Marois (2005) used a 12-alternative match-to-sample task (see similarly: Fougnie & Marois, 2007; Devue et al., 2009). And Most et al. (2005) asked an open-response follow-up: “If you did see something on the last trial that had not been present during the first two trials, what color was it? If you did not see something, please guess.” These questions are more difficult and to that extent less sensitive than binary forced-response/2afc questions of the sort we use in our own studies – a difference which may be critical in uncovering degraded perceptual sensitivity. For all these reasons, we believe our novel approach of using 2afc or forced-response questions combined with signal detection analysis is an important improvement on prior methods.

Previous studies of the related phenomenon of change blindness have investigated whether subjects who fail to detect changes nonetheless perform above chance in discrimination tasks concerning the changed object (Mitroff et al., 2002, 2004; Hollingworth & Henderson, 2002). However, only a handful of prior studies have explored the possibility that inattentionally blind subjects outperform chance in reporting or responding to features of IB stimuli (e.g., Schnuerch et al., 2016; see also Kreitz et al., 2020, Nobre et al., 2020). Moreover, a recent meta-analysis of this literature (Nobre et al., 2022) concluded that such work is problematic along a number of dimensions, including underpowered samples and evidence of publication bias that, when corrected for, eliminates effects revealed by earlier approaches. (These concerns hold in addition to our own worries about biased measures of performance.) The authors of this meta-analysis conclude with the following recommendation for future work: “We suggest that more evidence, particularly from well-powered pre-registered experiments, is needed before solid conclusions can be drawn regarding implicit processing during inattentional blindness” (Nobre et al., 2022). We see the present set of high-powered pre-registered studies as providing precisely this evidence, in ways that advance our understanding of IB considerably.

Our results also shed new light on evidence that inattentionally blind subjects process the unexpected stimuli they deny noticing. For example, in the electrophysiology literature, unexpected line patterns have been found to elicit the same Nd1 ERP component in both noticers and inattentionally blind subjects (Pitts et al., 2012). Likewise, preserved neural signatures of scene segmentation and perceptual inference (e.g., Kanizsa figures) have been found in inattentionally blind subjects (see respectively, Scholte et al., 2006, and Vandenbroucke et al., 2014), though interestingly not differential responses to meaningful words versus meaningless letter strings (Rees et al., 1999). Similarly, behavioral studies show that unattended stimuli can influence the accuracy and speed of inattentionally blind subjects’ judgments in a primary task (e.g., Moore & Egeth, 1997; replicated in Wood & Simons, 2019; Pugnaghi et al., 2020). Although some researchers have interpreted these results as implying a kind of subliminal processing of IB stimuli, our results just as easily raise an alternative (and perhaps more straightforward) explanation: that inattentionally blind subjects may retain a degree of awareness of these stimuli after all.

We acknowledge that above-chance performance in our experiments could be taken to reflect unconscious representations in a manner akin to orthodox interpretations of blindsight (Weiskrantz et al., 1974; Kolb & Braun, 1995; though see Phillips, 2021a; Morgan et al., 1997; and for more general skepticism, Newell & Shanks, 2023). However, in our view, explicit voluntary judgments of stimulus features (especially in neurotypical subjects) constitute prima facie evidence of conscious processing, and should be interpreted that way unless there is some compelling reason to favor an alternative (Snodgrass, 2002; Balsdon & Azzopardi, 2015; Heeks & Azzopardi, 2015; Phillips, 2021b). As a result, although we acknowledge the possibility that our data reflect unconscious processing, we think there are compelling reasons to interpret our results in terms of residual conscious vision in IB (although we note that this claim is tentative and secondary to our primary finding).

Evidence that inattentionally blind subjects process and are sensitive to the unexpected stimuli they deny noticing has been used to support so-called *inattentional amnesia* accounts of IB—the traditional rival to the orthodox interpretation of IB. On inattentional amnesia accounts, unattended objects and features are consciously perceived but not encoded so as to be available for later explicit report (Wolfe, 1999; Moore, 2001; though see Ward & Scholl, 2015; Hirschhorn et al., 2024). Our results are consistent with some degree of inattentional amnesia, and likewise, with what Block (2001) calls *inattentional inaccessibility*. (For a fuller discussion of these alternative hypotheses and IB more generally, see Wu, 2014.) However, our findings suggest that inattentional amnesia cannot be the whole story, since they reveal that some features of unexpected objects are available for later explicit report even in the group of subjects who deny noticing anything unusual.

### Visual awareness as graded

A further upshot of our findings is that they lend support to a more graded perspective on IB (in particular) and both perception and visual awareness (in general). This stands in contrast to the two interpretations that have dominated discussion of IB, both of which adopt a binary perspective. On the orthodox interpretation, inattention abolishes all perception and awareness; on the rival inattentional amnesia account, inattention abolishes all explicit encoding. Our data suggest the need to move beyond such binaries (cf. Cohen et al., 2023): Inattention degrades but does not eliminate perception and awareness – and likewise explicit encoding. This more nuanced approach has some kinship with what Simons (2000) calls *inattentional agnosia*. On this account, subjects who report not noticing may have some awareness of the unexpected object but fail to “encode the properties necessary to register that the item was something new, different, or noteworthy” (Most et al., 2005). Though aligned with the spirit of our view, this account still does not fully capture our results, since in our studies the group of non-noticing subjects could explicitly report features which would ordinarily suffice to mark the unexpected object as new or different (e.g., color, shape). Nonetheless, we agree that unattended stimuli are encoded in a partial or degraded way. Here we see a variety of promising options for future work to investigate. One is that unattended stimuli are only encoded as part of ensemble representations or summary scene statistics (Rosenholtz, 2011; Cohen et al., 2016). Another is that only certain basic “low-level” or “preattentive” features (see Wolfe & Utochkin, 2019 for discussion) can enter awareness without attention. A final possibility consistent with the present data is that observers can in principle perceive individual objects and higher-level features under inattention but that the precision of the corresponding representations is severely reduced. Our central aim here is to provide evidence that there is residual perceptual sensitivity to visual features for subjects who would ordinarily be classified as inattentionally blind. Further work is needed to characterize the exact nature of this perception, and the awareness (if any) which corresponds to it.

### Conclusion

Taken together, and after decades of inconclusive findings, the results of our five studies offer the strongest evidence so far of significant residual visual sensitivity across a range of visual features in IB. In other words, as a group, the inattentionally blind enjoy at least some degraded or partial sensitivity to the location, color and shape of stimuli which they report not noticing. Together with our finding that subjects collectively exhibit a systematically conservative bias in reporting their awareness, our results also call into question the orthodox interpretation of IB on which inattention entirely abolishes awareness, suggesting that a reconceptualization of inattentional blindness may be required. Indeed, perhaps ironically, inattentional blindness if anything provides evidence that awareness of certain features *survives* inattention. Our results highlight the critical value of assessing response bias and including objective measures of sensitivity in studying inattentional blindness and visual awareness. They also point to a broader rethinking of perception and consciousness as graded, rather than binary, phenomena.

## Methods

### Open Science Practices

All sample sizes, exclusion criteria, analyses and key experimental parameters reported here have been pre-registered. Data, analyses, stimuli and pre-registrations are publicly available at https://osf.io/fcrhu/. Readers can also experience all experiments for themselves at https://perceptionresearch.org/ib/.

### Experiment 1: Above-chance sensitivity to the location of unnoticed stimuli

#### Participants

500 adults were recruited from the online platform Prolific (for validation of the reliability of this subject pool, see Peer et al., 2017), with participation limited to US subjects. As described in our pre-registration, we reached this number by running batches of 100 subjects until a target number of 100 non-noticers (i.e., subjects answering “no” to the yes/no question about whether they noticed the unexpected stimulus) was reached. After excluding subjects who incorrectly reported which arm of the cross was longer on any of trials 1-3 and those who failed to provide a complete dataset, or failed a test for color vision (Ishihara color plate; see data archive), 374 subjects were included in the analysis. (All of these exclusion criteria were pre-registered.) This experiment and all others reported here were approved by the Homewood Institutional Review Board of Johns Hopkins University. All subjects provided informed consent and were compensated financially for their participation.

#### Stimuli and Procedure

As shown in Figure 2, Experiment 1 contained four trials, with the fourth trial differing from the first three in several ways. All trials took place in a display with dimensions 600 px x 600 px. Due to the nature of online experiments, we cannot be sure of the exact size or distance of stimuli as subjects actually experienced them (and so we give these figures in pixels); however, any differences in subjects’ monitors and/or display setups would have been constant across all trials of the experiment.

On trials 1-3, a fixation circle appeared in the center of the display; subjects pressed the spacebar when they were ready to begin the trial. The keypress was followed by a 1500 msec delay, after which a cross formed by two thin black lines (3 px thickness; 200 px x 140 px dimensions) appeared for 200 msec either 150 px above or below the fixation circle. The location of the cross (above or below), as well as its aspect ratio (vertical-longer or horizontal-longer) was randomly chosen on each of these trials. The cross then disappeared, followed by a 500 msec blank interval. Subjects were then asked which of the cross’s arms was longer (horizontal or vertical).

The fourth, critical trial proceeded the same way, except that simultaneous with the cross, a vertical red line (200 px long; 3 px thick; RGB(147,0,0)) appeared on one side of the display (10 px from the boundary of the display), also for 200 msec. Subjects were then asked the same cross arm length question as before.

Following this, two additional questions were asked, in the following order, each on its own display (such that subjects only saw the second question after answering the first).

Question 1

*Did you notice anything unusual on the last trial which wasn’t there on previous trials? (yes / no)*

Question 2

*In fact, the last trial you just saw contained one extra element — a vertical red line on either the left or the right side of the box. What side do you think it appeared on? If you don’t know, or don’t think it appeared on either side, take your best guess. (left / right)*

For all forced-choice and forced-response questions asked in all experiments reported here, subjects indicated their answers by clicking a radio button next to the text corresponding to their answer, and then submitted their responses by clicking a separate “Submit” button, a design aimed at eliminating motor-error responses.

#### Analysis and Results

As reported in the main text, 28.6% (107/374) of subjects responded “no” to Question 1; we refer to these subjects as “non-noticers”. These subjects are those who demonstrate inattentional blindness by conventional standards. However, 63.6% of non-noticers answered Question 2 correctly (the 2afc location task). This proportion was compared to chance responding (50%) with both frequentist and Bayesian null hypothesis tests, using the binom.test and proportionBF functions, respectively, from the BayesFactor package in R (Ihaka et al., 1996; Morey et al., 2015; all arguments used were defaults), which yielded a 95% CI = [0.54, 0.73], and a BF10 = 9.9.

SDT analyses began by calculating the number of hits, false alarms, present trials, and absent trials, and then applying the log-linear correction of adding 0.5 to all cells of the decision matrix (Hautus, 1995; see also Box 1). This is a standard correction to prevent an infinite *d′* in the event that either hits or false alarms are zero (this correction was applied to all experiments, though neither hits nor false alarms were ever zero), and simulations have shown that if anything, this correction *underestimates d′* (Hautus, 1995).

The signal detection theory measure of sensitivity (*d′*) for non-noticers’ performance on the 2afc location task was calculated as follows:

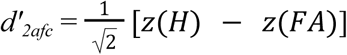

For our analysis, we (arbitrarily) considered trials where a stimulus was presented on the *left* to be “present” trials, and trials where a stimulus was presented on the *right* to be “absent” trials (*d′* will be identical regardless of which trial type is considered present/absent; *c* will be the same value with the opposite sign). With a hit rate and false alarm rate of 72.64% and 45.54% respectively (after log-linear adjustment), the resulting *d′_2afc_*= 0.51. Note that *d′* for this experiment was adjusted downward by a factor of 1/ 2 because 2afc tasks are theoretically easier than yes/no or forced-response tasks (Macmillan and Creelman, 2005). This formula for *d′_2afc_* (as well as those for *d′* and *criterion*, described below) assumes equal variance for signal and noise distributions (a standard assumption in SDT), but the analysis code provided allows for the calculation of SDT statistics when the variance of signal and noise distributions is unequal.

Because each subject in the experiment contributes just one trial, the signal detection metrics were calculated at the group level, and the standard error for each SDT calculation was estimated using methods described by Macmillan and Creelman (2005, pgs. 325-328; see also Kadlec, 1999). We estimated the variance for *d′* in this 2afc task using methods first described by Gourevitch and Galanter (1967), and re-described in Macmillan and Creelman (2005), equations 13.5 and 13.7. First, equation 13.5 demonstrates how ϕ (a function which converts z-scores into probabilities) can be computed for the hit rate and false alarm rates:

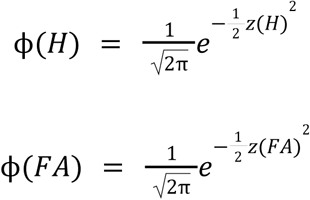

With *ϕ*(H) and *ϕ*(FA) computed, we then estimated the variance of *d′* in this 2afc task using equation 13.7:

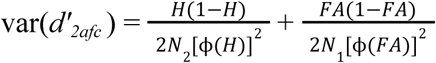

where N_2_ is the number of present trials, and N_1_ is the number of absent trials. (For information about how the variance is affected by sample size using different methods and in different tasks, see Macmillan and Creelman Table 13.2 and 13.3; our sample sizes are more than sufficient to expect this variance estimation to be accurate to the hundredth decimal place).

Finally, we computed a confidence interval around *d′_2afc_* using standard methods: The result is 95% CI = [0.16, 0.85], suggesting performance in the non-noticing group was significantly above chance.

### Experiment 2: Above-chance sensitivity to the color of unnoticed stimuli

#### Participants

1700 adults were recruited from Prolific, collected in batches of 100 subjects until a target number of 100 non-noticers was reached. After excluding subjects who incorrectly reported which arm of the cross was longer on any of trials 1-3 and those who failed to provide a complete dataset, or failed a test for color vision, 1261 subjects were included in the analysis.

#### Stimuli and Procedure

The fourth, critical trial proceeded the same way as Experiment 1, except that the extra vertical line that appeared simultaneous with the cross was either red (RGB(147,0,0)) or blue (RGB(0,0,136)), with the color and location of the line randomized across subjects.

Following the presentation of Trial 4, subjects were asked which cross arm was longer, and then the same traditional IB question as in Experiment 1 (Question 1), followed by:

Question 2

*The last trial you just saw contained one extra element — a vertical line on one side of the box. What color was the extra line? If you don’t know, or don’t think any line appeared, take your best guess. (red / blue)*

To reduce uncertainty about what color “red” and “blue” referred to, the text for each color option was printed in the red and blue color used for the IB stimuli in the different conditions (RGB(147,0,0), RGB(0,0,136)).

#### Analyses and Results

As reported in the main text, 27.7% of subjects shown an additional stimulus responded “no” to Question 1 (i.e., demonstrated inattentional blindness by conventional standards). However, 58.5% of these non-noticers answered correctly on Question 2 (95% CI = [51.95%, 64.93%]; BF10 = 4.54).

In Experiment 2, the follow-up color discrimination was a two-alternative forced-response design, and so for signal detection analyses, sensitivity was calculated without the 1/sqrt(2) adjustment, such that

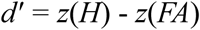

resulting in a *d′* = 0.38. We estimated the variance of *d′* for this one-interval forced-response task using similar methods to those described for Experiment 1, with one minor change to the variance equation to account for this being a forced-response task (equation 13.4 in Macmillan and Creelman, 2005):

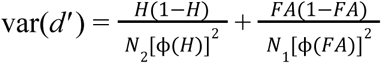

The only difference between equation 13.4 and equation 13.7 is that the latter includes a factor of 2 in the denominator of both terms, which accounts for 2afc tasks theoretically being easier than yes/no tasks.

The procedure for significance testing comparing *d′* to chance (*d′* = 0) was identical to Experiment 1 Methods, and the two-sided frequentist binomial probability test yielded a 95% confidence interval of [0.03, 0.73]. As stated in the main text, subjects demonstrated a significant bias to respond “blue,” which we measured by calculating subjects’ criterion on the red/blue question as,

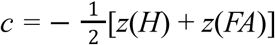

resulting in a positive (conservative) criterion value of *c* = 0.67. The 95% confidence interval around the criterion estimate is [0.49, 0.84]. Since this interval does not contain zero, this represents a statistically significant bias. For a more intuitive understanding of the bias, 78.42% of subjects shown *no IB stimulus* guessed that the stimulus was blue.

Lastly, an important contribution of this work is the inclusion of absent trials, which enable us to compute response bias (*c*) for the traditional IB question, “Did you notice anything unusual on the last trial that wasn’t there on previous trials?” As predicted, we found that, as a group, subjects were conservative in reporting their awareness of the IB stimulus (*c* = 0.31, 95% CI = [0.22, 0.41]), suggesting that subjects in inattentional blindness experiments may systematically underreport their awareness of unexpected stimuli.

### Experiment 3: Above-chance sensitivity even in highly confident non-noticers

#### Participants

7,000 subjects were recruited from Prolific, and data were collected in batches until we reached 200 non-noticers who reported being “highly confident” in their yes/no response. After excluding duplicate data files, subjects who incorrectly reported which arm of the cross was longer on any of trials 1-3, those who failed to provide a complete dataset, or failed a test for color vision, 5296 subjects were included in the analysis.

#### Stimuli and Procedure

This experiment explored whether sensitivity to the visual features of an IB stimulus varies as a function of subjects’ confidence in their responses to the traditional IB question (i.e., whether or not they noticed anything unusual on the last trial), and more specifically whether even highly confident non-noticers would show such sensitivity. Experiment 3 was identical to Experiment 1, except that after subjects answered the traditional yes/no question (which was the same as Question 1 in Experiment 1) and the follow-up left/right question (which was the same as Question 2 in Experiment 2), they also rated their confidence in each of those responses:

Question 1

*Did you notice anything unusual on the last trial which wasn’t there on previous trials? (yes / no)*

Question 2 (Confidence rating for Question 1)

*How confident are you in your answer? (Four-point scale: 0 = Not at all confident – 3 = Highly confident)*

Question 3

*The last trial you just saw contained one extra element - a vertical red line on either the left or the right side of the box. What side do you think it appeared on? If you don’t know, or don’t think it appeared on either side, just take your best guess. (left / right)*

Question 4 (confidence rating for Question 3; not shown in Figure 3)

*How confident are you in your answer? (Four-point scale: 0 = Not at all confident – 3 = Highly confident)*

#### Analyses and Results

As reported in the main text, 30.85% of subjects shown an additional stimulus responded “no” to Question 1 (i.e., demonstrated inattentional blindness by conventional standards)

As with Experiments 1 and 2, we were interested in whether subjects who responded “no” to the traditional IB question would nevertheless perform above-chance on subsequent discrimination questions about the IB stimulus. Beyond our general interest in all subjects who answered “no,” we were most interested in whether the non-noticers who reported high confidence in their answer still performed above-chance on the 2afc question. We compared the performance of those observers to chance (*d′* = 0) using the methods described above, and found that even the most confident non-noticers—i.e., those who reported being “highly confident” that they did not notice anything unusual, rating their confidence at 3 on a scale from 0 to 3—collectively demonstrated significantly above-chance sensitivity to the location of the IB stimulus: *d*′_2afc_ = 0.34; 95% CI = [0.08, 0.60]. Importantly, this group of subjects is minimally powered, with just 204 subjects, adding force to the argument that there is meaningful sensitivity amongst highly confident non-noticers. The bin of second-most interest is that of the moderately confident non-noticers (those responding with a confidence rating of 2 on a scale from 0 to 3). Sensitivity in this group was above zero (*d*′_2afc_ = 0.05), but not significantly so (95% CI = [-0.10, 0.20]).

Figure 3e depicts the uncorrected results of null hypothesis significance tests comparing the *d*′_2afc_ estimate for each confidence rating bin to chance (*d*′_2afc_ = 0), but essentially the same pattern of significant results is obtained when the Holm-Bonferroni correction for multiple comparisons is applied (*p* = .034 for the No-3 bin, and *p* = 0.09 for the Yes-0 bin–though note this bin only contains 25 subjects).

### Experiment 4: Above-chance sensitivity in a sustained inattentional blindness task

#### Participants

1500 subjects were recruited from Prolific, and data were collected in batches until we reached 100 non-noticers who answered the color discrimination question first, 100 non-noticers who answered the shape discrimination question first, and 100 non-noticers who answered the location discrimination question first (more on these three discrimination questions below).

After exclusions, 1278 subjects including 417 non-noticers remained in the analysis for Experiment 4. The pre-registered exclusion criteria for this experiment were the same as those reported by Wood & Simons (2017). Subjects were excluded if: (i) their reported bounces for either of the first two trials erred by more than 50% in either direction from the true number of bounces of their attended objects on that trial, (ii) they failed to contribute a complete dataset, (iii) they reported problems with experimental playback, such as stuttering, freezing or another issue specified in a free-response, or (iv) an observer managed to submit or run the study twice (evidenced by two files sharing the same Prolific ID), in which case we excluded their second run from the analysis.

#### Stimuli and Procedure

In Experiment 4, subjects participated in a sustained inattentional blindness task modified slightly from experimental code published by Wood & Simons (2017). All subjects completed three trials of a dynamic, multiple object tracking task containing black squares and white squares. At the beginning of the experiment, each subject was told whether they should attend to the black or white squares. At the beginning of each trial, subjects were instructed to fixate on a small blue square (11 px x 11 px; RGB(0,0,255)) in the center of a gray (RGB(127,127,127)) rectangle (666 px x 546 px), and were told that their task was to count the number of times the squares of their attended color bounced off of the walls of the gray rectangle. Each trial lasted approximately 17 seconds, and each subset of black/white squares bounced an average of 28 times. After the trial ended, subjects were asked to report how many times the squares of their attended color bounced off of the walls of the rectangle. Of the 222 excluded subjects, 185 were excluded for bounce reports that erred by more than 50% in either direction of the actual number of bounces.

The critical trial was Trial 3. In this trial, 3/4 of subjects (Present condition) were shown an unexpected shape (a circle or a triangle, which was either black or white). This unexpected object entered the display on either the left or the right 5 seconds after the trial began, and moved either upward or downward until it exited the other side of the display (thus, there were 2 color x 2 shape x 2 side x 2 motion direction options for IB stimuli, randomly chosen for each subject). For 1/4 of subjects, no additional object was shown on Trial 3 (Absent condition).

Regardless of condition, at the end of the critical trial, subjects were again asked to report the number of bounces, followed by four additional questions:

Question 1

*Did you notice something on the last trial that did not appear on previous trials? (yes/no)*

After answering Question 1 (the standard IB question), subjects were told (again regardless of condition): *An extra object may have appeared on that last trial. If you saw it, please tell us its color, shape, and whether it appeared on the left or the right side of this gray box*. *If you didn’t see the extra object, please guess. We’ll ask about* [whichever discrimination question was randomly chosen to be asked first] *first*.

Questions 1-3 (presented in random order for each subject)

*The new object was… (black/white)*

*The new object was a… (circle/triangle) The new object was on the… (left/right)*

Note that Question 1 (the traditional IB question) differs slightly in Experiments 4 and 5 from the question wording in Experiments 1-3 because we aimed to cleave as closely as possible to Wood & Simons (2017) and other inattentional blindness experiments using similar paradigms.

#### Analyses and Results

As reported in the main text, 57.32% of subjects shown an additional stimulus on the critical trial answered “no” to the traditional IB question (Question 1). For absent trials, 6.31% of subjects answered “yes” to the traditional IB question when no additional stimulus appeared; this is the false alarm rate, which can be used (along with the hit rate) to estimate subjects’ bias in responding to the traditional IB question. In Experiment 4, as in Experiment 2, we found that, as a group, subjects answered the traditional IB question using a conservative criterion (*c* = 0.85, 95% CI = [0.73, 0.97]), suggesting they may be underreporting their awareness of IB stimuli.

Experiment 4 asked subjects to perform 3 discrimination tasks following their response to Question 1, with the order randomized for each subject. The color and shape discriminations are one-interval forced response tasks, and so *d′* was calculated for these two questions without the 2afc correction. We found that non-noticers collectively performed significantly above chance on both the color (*d′* = 0.82, 95% CI = [0.61, 1.04]) and shape (*d′* = 0.21, 95% CI = [0.005, 0.42]) discriminations. In addition to the issue identified below, the fact that this result was marginal further motivated Experiment 5, in which we substantially increased the number of subjects recruited in order to reduce the size of the confidence intervals around *d′* estimates of discrimination sensitivity by 50%. For the location discrimination (left/right), the 2afc correction to *d′* was applied, and non-noticers’ performance was trending but was not above-chance (*d′*_2afc_ = 0.07, 95% CI = [-0.08, 0.22]).

Because the order of the discrimination questions varied by subject, we pre-registered an analysis specifying that we would also derive sensitivity and bias for subjects who answered a given discrimination question first. With these subsets of non-noticers (roughly one-third the original sample size), confidence intervals are much larger, and although color discrimination remained significantly above-chance (*d′* = 0.78, 95% CI = [0.41, 1.16]; N = 189), shape discrimination was no longer significant (*d′* = 0.15, 95% CI = [-0.20, 0.50]; N = 195) and location (*d′*_2afc_ = -0.17, 95% CI = [-0.44, 0.10]; N = 176) was again not significant.

Finally, in a pre-registered analysis breaking down the inattentional blindness rate by the color of the attended stimuli, we found that subjects were roughly 3x more likely to report noticing a color-congruent than a color-incongruent IB stimulus (86.57% vs. 28.71%). This interacted with subjects’ responses on the color discrimination task, with subjects shown an IB stimulus demonstrating a significant bias to say that the stimulus that appeared was the opposite color to the colored squares they attended (84.25% of subjects who attended to white squares answered black; 75.09% of subjects who attended to black squares answered white). In order to get a better estimate of sensitivity to color in this task, we pilot tested a new pair of IB stimulus colors in Experiment 5 to equate the guessing rate on absent trials.

### Experiment 5: Replicating above-chance sensitivity in a sustained inattentional blindness task

#### Participants

To ensure a large enough sample without overlap with previous experiments using this paradigm, Experiment 5 recruited Prolific subjects not only from the USA but also from Canada, the United Kingdom, and Australia. In order to reduce the size of the confidence intervals around the sensitivity estimates relative to Experiment 4, we collected data until we reached at least 2,200 subjects who reported not noticing the unexpected stimulus. Exclusion criteria were identical to Experiment 4. After exclusions, 10,830 subjects in total, including 2,339 non-noticers, were included in the analysis. (To our knowledge, this made Experiment 5 the largest single inattentional blindness sample ever collected.)

#### Stimuli and Procedure

Experiment 5 repeated the design of Experiment 4, with two key differences: (1) Every subject was assigned to attend to the white squares in the primary task, (2) the IB stimuli were either orange or green—two colors that as a pair produced equal IB rates in pilot testing. Again, subjects were asked the traditional IB question (yes/no) followed by three discrimination questions (color, shape, and location) in random order for each subject. The change to the colors of the IB stimuli of course meant that the color question and options were changed to read:

*The new object was… (color 1/color 2; with the text printed in the given color)*

#### Analyses and Results

As reported in the main text, with the congruency effects mitigated and despite the highly salient color of the orange/green IB stimuli, the pattern of results matched that of Experiment 4: Analysis of responses to the traditional yes/no question once again revealed that subjects were collectively biased to respond “no” (*c* = 0.45, 95% CI = [0.41, 0.49]), with the hit rate (subjects in the Present condition who responded “yes”) being 71.01% (100% - the IB rate) and the false alarm rate (subjects in the Absent condition who responded “yes”) being 7.24% (see a more detailed breakdown of the SDT analysis for this experiment in Box 1). Thus, even when a highly salient, moving stimulus entered the display suddenly and remained on screen for multiple seconds, subjects were hesitant to report noticing anything unusual.

Non-noticers’ performance on the three discrimination questions was also consistent with the pattern of results in Experiment 4, with non-noticers collectively performing significantly above-chance in discriminating both color (*d′* = 0.12, 95% CI = [0.02, 0.23]) and shape (*d′* = 0.23, 95% CI = [0.13, 0.33]), but not location (*d′_2afc_* = 0.03, 95% CI = [-0.04, 0.11]). Biases in the discrimination question responses among subjects shown no IB stimulus on the critical trial (the Absent condition) were minimal, confirming that the congruency effect (interaction between the IB rate and subjects’ choices on the color discrimination question) was mitigated in Experiment 5: For subjects shown no unexpected stimulus on the critical trial (the Absent condition), 52.95% guessed that the extra object had been orange, and 47.05% guessed that it had been green. On the shape question, 51.43% of subjects guessed that the extra object had been a triangle, and 48.57% guessed that it had been a circle. Finally, on the location question, 54.87% of subjects guessed that the extra object had been on the left side of the display, and 45.13% guessed that it had been on the right side of the display.

## Footnotes

1 See our Supplementary Material for reason to think that this may actually *underestimate* subjects’ true performance, which may be nearly double this figure.

